# NuCDB: A databank of nucleic acids circular dichroism spectra

**DOI:** 10.1101/2024.04.19.590190

**Authors:** Uttam Das, Murali Aadhitya Magateshvaren Saras, Thenmalarchelvi Rathinavelan

## Abstract

Nucleic acids take a variety of secondary structures depending on the sequence and environmental conditions. Circular dichroism (CD) spectroscopy quickly provides the secondary structural information of nucleic acids. **Nu**cleic acid **C**ircular-dichroism **D**ata**B**ank (NuCDB) https://project.iith.ac.in/nucdb/, a repository of nucleic acid secondary structure CD spectra published during 1964-2012, is created. Besides acting as the repository for the nucleic acids CD spectra, **NuCDB** also has the facility to upload recently published CD spectra to keep the repository up-to-date. This repository provides the sequence-structure-environmental relationship of different nucleic acid fragments in one platform. Until today, CD is used for studying the secondary structure of smaller nucleic acid fragments. Since different parts of the genome and transcriptome of an organism have combinations of various secondary structures and play a crucial role in regulating the biological processes, the CD spectra of longer nucleic acid sequences would be more realistic. Thus, this bioinformatics repository would be helpful in training machine learning models to predict the presence of multiple secondary structures in a given CD spectra.

## INTRODUCTION

Circular dichroism (CD) spectroscopy is a technique that employs the difference in the absorbance of right- and left-circularly polarized light by any substance. An inherently asymmetric chromophore that absorbs radiation in its ground state at ambient temperature produces a CD spectrum. The CD spectra are produced when a plane-polarized electromagnetic wave which encompasses two circularly polarized vectors (right-handed and left-handed each) of equal intensity traverses through an asymmetric chromophore. An asymmetric chromophore absorbs these two components of plane-polarized compounds at different extents, resulting in an elliptical vector describing the ratio of the major and minor axes that determines the ellipticity, the phenomenon known as optical activity.

CD spectroscopy is extensively used for studying the secondary structure of biopolymers, specifically proteins and nucleic acids. It is relatively easy to identify the secondary structure of proteins from the CD spectra on account of folding constraints and the restricted possibilities, which include alpha helix, beta sheet, and random coil. Differentiating the CD spectra of nucleic acids can be challenging due to the diverse secondary structures observed. Several investigations have been carried out to predict the secondary structure of proteins and nucleic acids using machine learning algorithms from the CD spectra(1-6). A well-organized repository of CD spectra of proteins and nucleic acids is essential to train the machine learning models. Considering this point, efforts have been made to create a CD spectral repository of proteins, namely, the protein circular dichroism databank (PCDDB)(6, 7). Here, we have established a database called NuCDB (**<u>Nu</u>**cleic acids **<u>C</u>**ircular **<u>D</u>**ichroism Data**<u>b</u>**ank) by collecting the nucleic acids (RNA and DNA) secondary structures through a comprehensive, systematic literature survey since the publication of the first nucleic acid CD spectra in 1964(8).

NuCDB (https://project.iith.ac.in/nucdb/) is currently a manually curated repository of 501 CD spectra of 17 nucleic acid secondary structures that has information about their spectral characteristics in the wavelength region of 190 nm to 320 nm. Each CD spectral data record has information about the spectral image and the concomitant sequence, environmental conditions, and the link to the published article. In short, NuCDB provides information about the sequence, secondary structure, and environmental conditions of nucleic acids. Thus, it can serve as a valuable resource to train machine learning models for predicting higher-order nucleic acid structures present in longer nucleic acid fragments. Additionally, it can be valuable for monitoring the alterations in the CD spectral peak during the process of titration with small molecule ligands and proteins. Since NuCDB is accessible free of cost with a substantial collection of DNA, RNA, and their hybrid CD spectral data along with the sequence and experimental conditions, it can help explore the structural complexity of nucleic acids more effectively.

## METHODS

### Data source and collection

The circular dichroism spectra corresponding to 17 types (501 data) of secondary structures of nucleic acids were fetched from the literature by searching in PubMed with the help of NCBI entrez programming utilities using the keywords “circular dichroism” “nucleic acid” or “CD spectra” and “DNA” or “CD spectra” and “RNA”. Following the retrieval of articles from PubMed, they were organized chronologically based on the year using NCBI filters. Subsequently, the CD spectral data reported in each article was manually examined. The CD spectral image (data) exclusively belonging to nucleic acids was specifically considered for creating this repository. Note that the CD spectra corresponding to the titration of nucleic acids with small molecule ligands are not considered. When more than one CD spectra are reported in an article, only a representative spectral image was downloaded and used in creating the dataset. Additionally, the experimental conditions, such as sample concentration, pH, and buffer details, along with the sequence information, were collected.

## Data Preparation

### Conversion of spectral images into numerical values

The downloaded CD spectral picture was manually transformed into a Comma Separated Values (CSV) format text file using WebPlotDigitizer 4.69(9). The CSV files contain data on the relationship between wavelength (nm) and ellipticity (θ in millidegrees) or molar ellipticity ([θ] in degree.cm2/dmol). Subsequently, they were used for plotting the CD spectral graphs (dark-red colored) and the corresponding images were created. Overall, 501 CD spectra published between the years 1964-2012 were stored both in CSV and image formats (**Figure 1**). This dataset is being updated to cover the published nucleic acid CD spectra until 2023.

**Figure 1.**
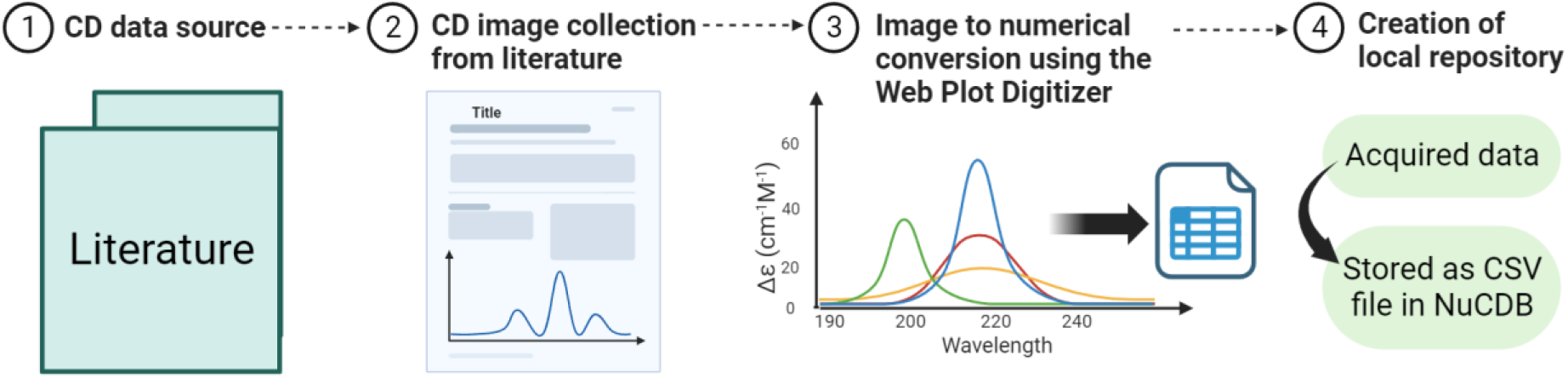
Schematic illustration of the data collection, processing, and creation of local repository.

## FUNCTIONALITY OF THE DATABASE

The dataset can be accessed freely through the provided link: https://project.iith.ac.in/nucdb/databank.html. The primary page of the databank formats (**Figure 2**) has four columns that provide summary of each CD spectrum. Each CD spectrum is assigned with a distinct reference ID, which is displayed in the first column. Note that the reference ID starts with a five-letter code which refers to different nucleic acids secondary structure (for example, APLQX stands for antiparallel (APL) quadruplex (QX)) followed by a serialized four-digit number. In addition to the reference ID, information regarding the nucleic acid type, secondary structure, and sequence is provided. Detailed information about each CD spectrum, such as the sample concentration, spectral wavelength range, molar ellipticity or ellpiticity values (as applicable), PDB ID (as applicable), experimental conditions and DOI information can be browsed by clicking on the reference ID link. Furthermore, the created spectral graph will also be displayed. Each reference ID is linked with a corresponding CSV file which has 2 column spectral data (wavelength (in nm) versus molar ellipticity or ellipticity values). This data can be used for plotting the spectral graph in the Cartesian space. All data described here can be downloaded from the webpage (https://project.iith.ac.in/nucdb/).

**Figure 2.**
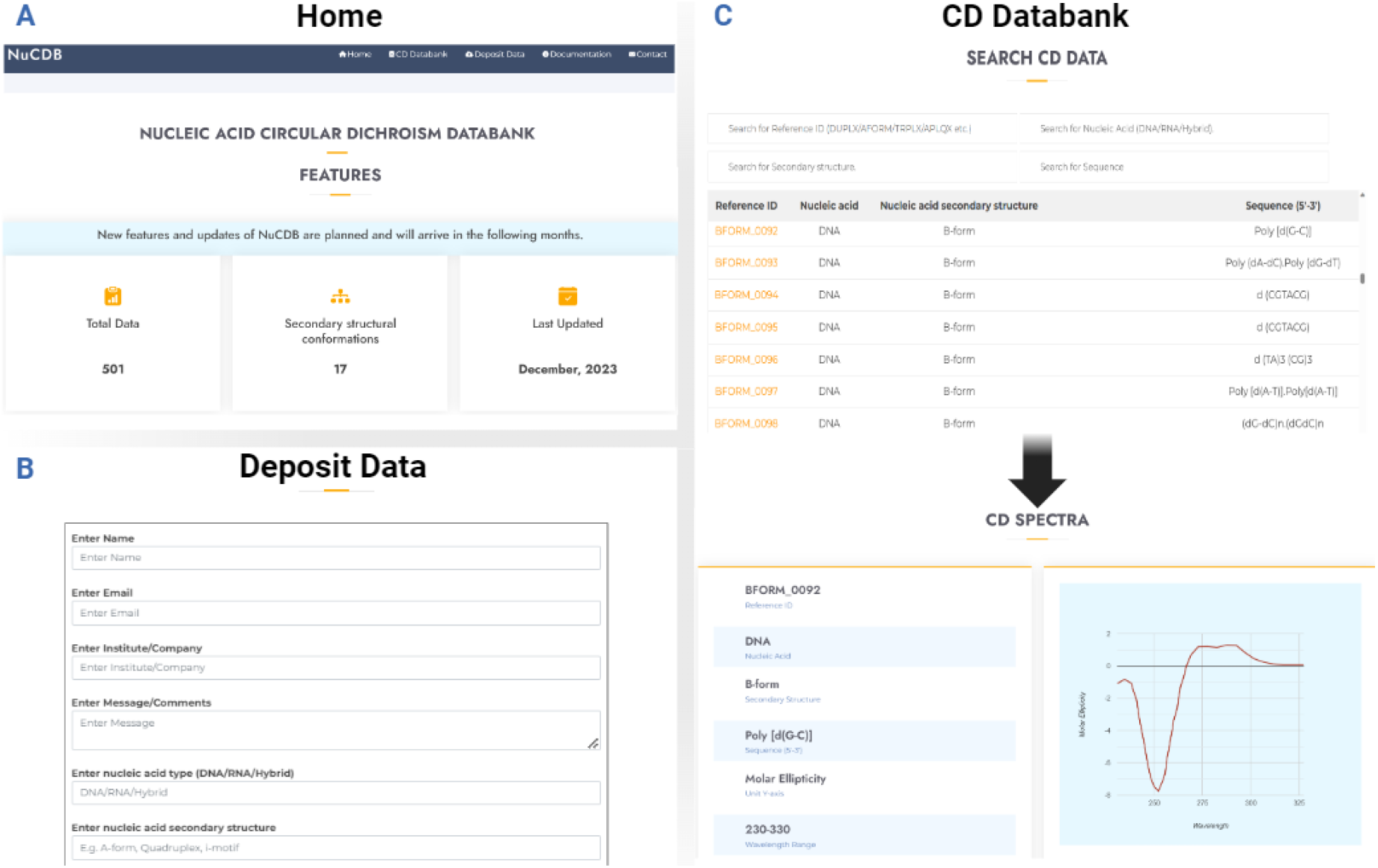
Snapshots showing the web interface of NuCDB: **(A)** Home Page, **(B)** depositing CD data, and **(C)** accessing CD data.

## TECHNICAL DETAILS

### Data storage

The complete list of metadata corresponding to 501 CD dataset belonging to 17 different types of secondary structures of nucleic acids is stored in a primary CSV (downloadable). Further, the numerical values (Wavelength vs Molar Ellipticity/Ellipticity values) of the 501 CD spectra are stored in individual CSV files. These spectral data files are available as tar-compressed and zip-compressed for download.

### Data validation

The CD spectral dataset was manually extracted from the sources referenced along with each dataset. The first author verified the classification of the dataset and the design of the data record, and all three authors agreed on the data entries and categorization. Since the CD spectral images are transformed into numerical data using WebPlotDigitizer software, the generated spectral points do not represent the actual experimental data points as the software generates the data points to mimic the shape of the spectra. Notably, such a transformation would not affect the characteristic features of the CD spectra corresponding to different nucleic acid secondary structures. This can be seen from random spectra taken for 17 different nucleic acid secondary structures, wherein the signature peaks of the original spectra are retained in the generated spectra (**Figure 3**).

**Figure 3.**
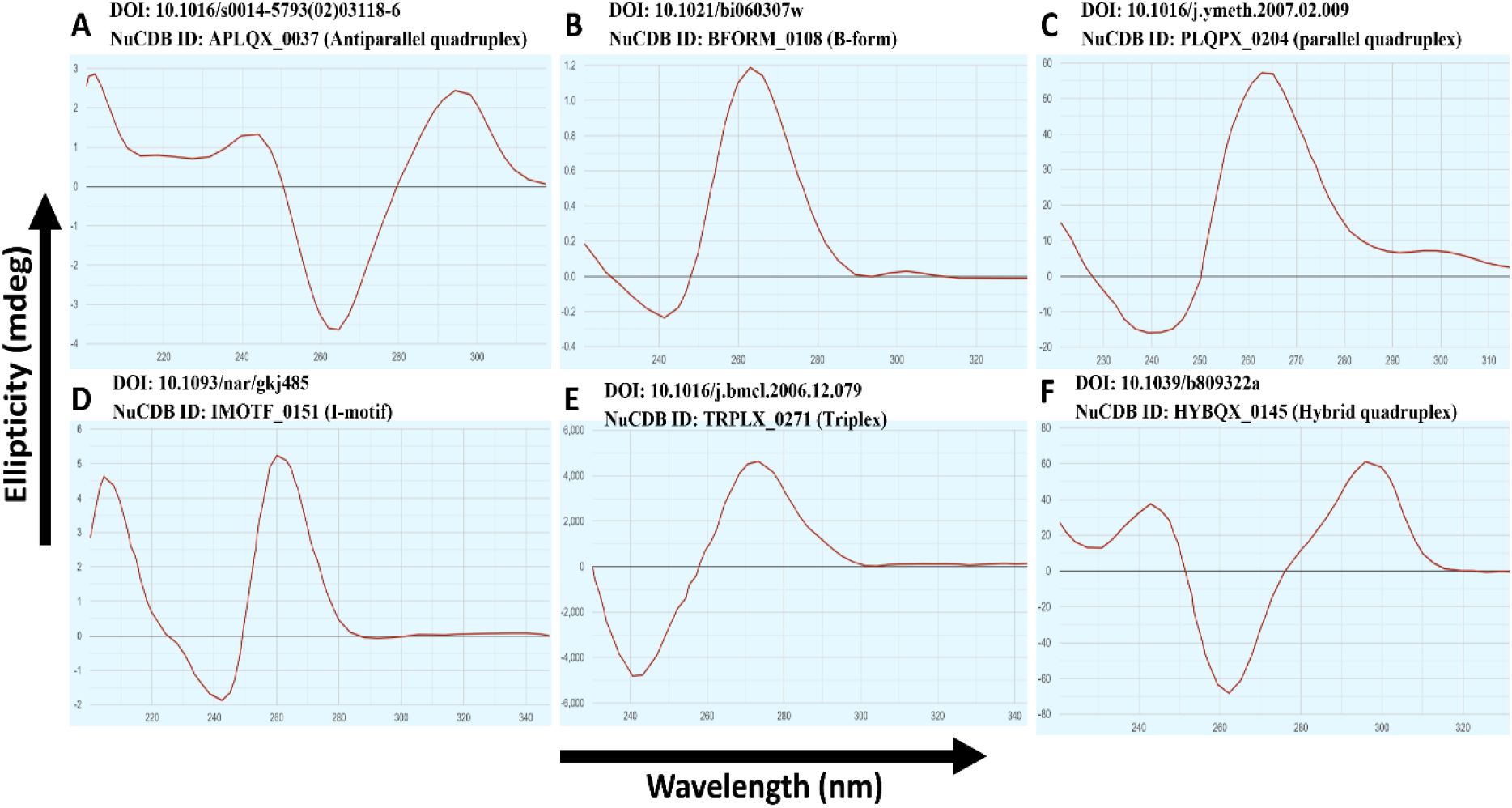
Illustrating few examples of the processed CD spectra fetched from its original source (X-axis and Y-axis represent wavelength (nm) and ellipticity (mdeg) respectively). The corresponding source (DOI), NuCDB ID and secondary structure are included alongside. (10-15).

## User notes

The CD spectral data of nucleic acids are available on a web platform (https://project.iith.ac.in/nucdb/), which will be updated with new outputs. A visual representation of each data can be accessed under the “CD Databank” menu (https://project.iith.ac.in/nucdb/databank.html). Selecting the molecule type and a corresponding secondary structure **(Figure 4; Steps 1-2-3)** assists in filtering the results. In addition, users have the option to search for a specific spectrum by inputting either a reference ID or sequence information. While the sequence and secondary structural information are directly displayed in a table format, the CD spectral image can be accessed by clicking the link under the “Reference ID” column. Further, users may also submit their data via this page along with the digital object identifier (DOI) of the corresponding published article **(Figure 4; Steps 1-4-5)**.

**Figure 4.**
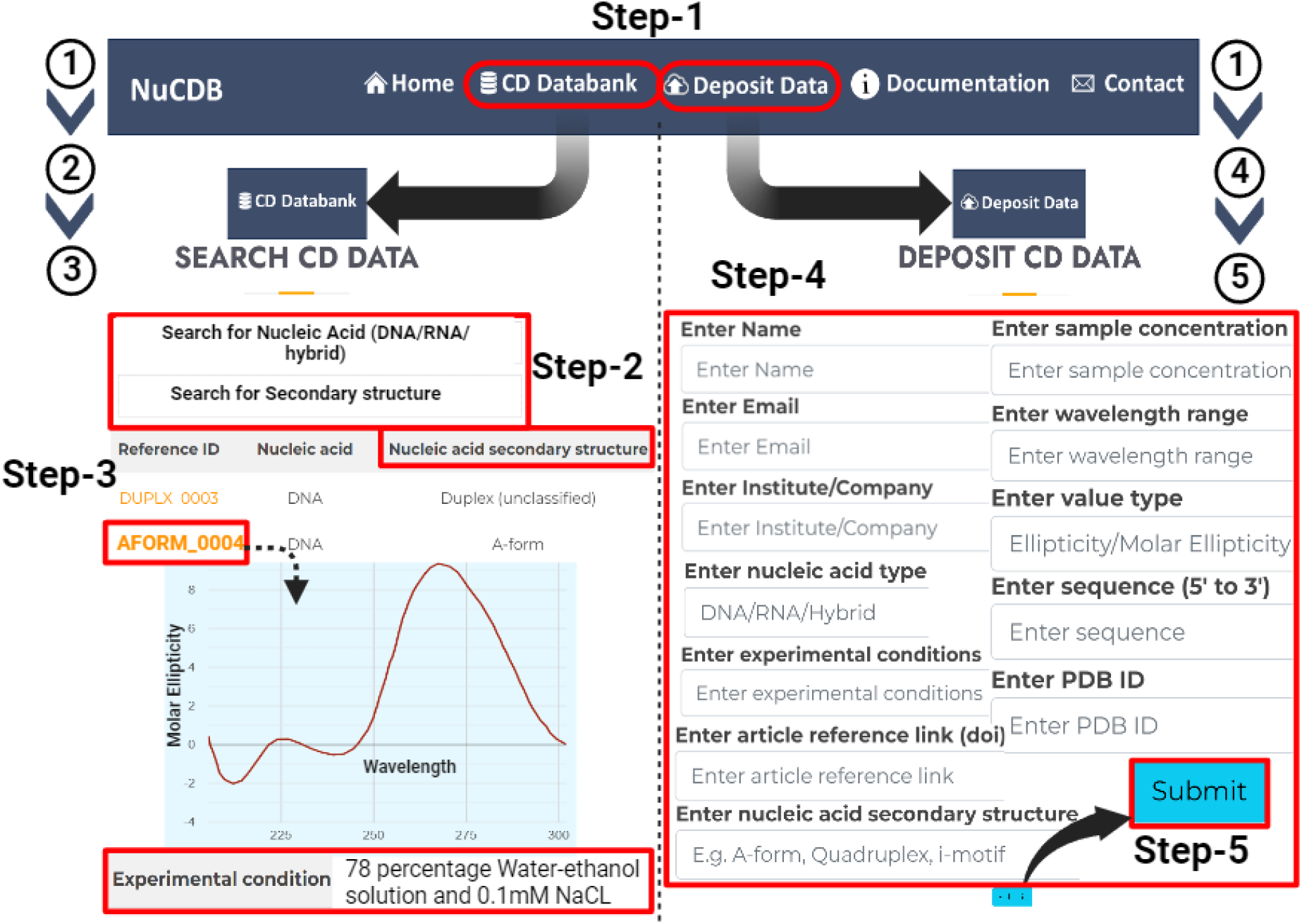
CD spectral data retrieval and deposition in NuCDB. (Left) Accessing the dataset from NuCDB illustrated with a representative data record (Steps 1-2-3). (Right) Steps (1-4-5) involved in new spectral deposition.

## SUMMARY AND FUTURE DIRECTION

NuCDB stands as a user-friendly database containing 501 manually curated circular dichroism spectral data linked to the secondary structures of 17 nucleic acids. It presents detailed information about the experimental conditions for each CD spectrum, accompanied by a visual representation of the spectral image. Moreover, users have the option to download the corresponding spectral data in CSV format, facilitating its application in various areas, such as training machine learning models. Although the database currently covers the CD spectral data up to 2012, NuCDB will soon be updated until 2023. Recognizing the importance of maintaining an updated dataset, NuCDB also enables users to upload published CD spectral datasets.

## AUTHOR CONTRIBUTION

UD collected and analyzed the CD spectral data. MAMS developed the web server. TR, UD, and MAMS wrote the manuscript. TR designed and supervised the entire project.

## ACKNOWLEDGEMENTS

The authors thank Sathyaseelan Chakkarai for his valuable suggestions during the early stage of the project. UD thanks UGC and MAMS thanks MoE for fellowships. The Computational Center, IIT Hyderabad, is acknowledged for its support.

